# PEEK: integrated primer design for silent-mutation and restriction-site engineering, validated in silico and by whole-plasmid sequencing

**DOI:** 10.64898/2026.10.29.762953

**Authors:** Kevin Ghobadi, Nathan Kura, John Warrington, Nathan Singh

## Abstract

**Motivation:** Silent introduction of restriction sites, direct site insertion, and verification of the result are routine operations in molecular cloning, yet they are typically distributed across separate tools built on inconsistent thermodynamics. Primer design is itself mature and heavily validated; the gap is not accuracy in principle but integration, together with the absence of any explicit account of why one synonymous edit should be preferred over another. Because melting temperature (Tm) gates which primers a tool will accept, any unified tool must first be shown to compute Tm correctly against an external standard, and then shown to produce the edits it proposes.

**Results:** PEEK, a web-based toolkit, unifies silent-mutation discovery, restriction-site insertion, and amplification and screening primer design built on a single thermodynamic model of primer melting temperature (Tm). PEEK’s silent-mutation engine identifies the smallest set of synonymous changes required to form a restriction site at a specific position, and ranks the options by codon usage, local mRNA folding, and edit size. Predicted melting temperatures closely matched those of the NEB Q5 Tm Calculator across 17 primers of varying length and GC content (Pearson R^2^ = 0.9995, mean absolute error 1.32 °C) without any fitted correction, and annealing-temperature (Ta) recommendations exactly matched NEB’s rules. In independent trials, a total of 15 silent-mutation designs were translationally silent and contained the intended restriction site. Experimental validation was performed by installing a KpnI site into a 9.9 kb lentiviral CAR plasmid using both a silent-mutation primer and an insertion primer. Whole-plasmid sequencing confirmed that each primer introduced exactly the designed changes, with no off-target differences across the construct and no change to the encoded protein.

**Availability and implementation:** PEEK runs at https://peekbiology.com and requires no registration. Source code, documentation, and test data are available at https://github.com/nkura-arch/PEEK-Protein-Engineering-Efficiency-Kit. Implemented in Python 3.11 with Flask, Biopython, and ViennaRNA.

## INTRODUCTION

Plasmid modifications are typically computed in small increments. Adding a restriction site to an existing plasmid construct additionally requires researchers to choose between two strategies. The first is site-directed silent mutagenesis, which introduces synonymous substitutions that create a restriction enzyme recognition sequence in a desired region without altering the encoded protein sequence itself. The second method, site-directed insertion mutagenesis, simply inserts the restriction site directly as a new sequence into the genetic code, altering the protein sequence as a byproduct. Both methods require primers to facilitate the Polymerase Chain Reaction (PCR) with melting temperatures falling within a range the polymerase can tolerate. Normally, they are verified after the experiment by amplifying the edited region the PCR targets and comparing it against the parent plasmid construct.

Modern tools exist for each of these steps. Primer3 (1, 2) acts as a general-purpose primer selection tool and has been consistently revised over its lifespan, including a major revision of its thermodynamic tables that was motivated in part by the inadequacy of its original Tm model (1). Primer-BLAST (3) adds to the functionality of Primer3 by searching for candidate primers against sequence databases, which improves efficiency. These commercial packages provide simple restriction-site mutation utilities; however, they work under proprietary algorithms that fail to fuse the total scope of primer design tools. What’s missing is consequently not a broad capability but rather the integration of these tools; a researcher designing a silent restriction site, the insertion alternative, and the screening primers for comparison purposes is currently required to manually move between tools with Tm models that disagree with each other, along with the polymerase manufacturer’s own Tm calculator.

In regard to silent mutations, codon optimization tools currently work to rewrite coding sequences in order to maximize expression of a target within a host where restriction sites are treated as constraints to avoid. Tools that address the inverse problem of introducing such restriction sites silently are much more sparse. SDM-Assist (4) is an application that designs site-directed mutagenesis primers that introduce silent restriction sites as a marker on bacterial clones. It ranks candidate primers with a weighted score across seven primer parameters, including GC content, melting temperature, number of nucleotide changes, etc. The authors of the application themselves note that unintended mutations (even silent ones) may influence expression efficiency of the primer depending on the organism or expression system. Other less effective tools even enumerate the synonymous changes without any ranking system whatsoever. This poses a large risk to the ability of modern primer design applications to effectively provide controlled, ranked mutations.

The PEEK silent mutation engine battles this issue by scoring the edit itself rather than the sole primers. It searches for the smallest local set of synonymous substitutions that introduce the requested restriction site at a user-input position. It then ranks the candidate substitutions by a function that accounts for the codon adaptation index, rare-codon clustering, local mRNA folding energy, and edit size, reporting each term individually and giving each a relative weight. We make no direct claim that this score predicts protein yield, as a multitude of external factors may impact this: each component’s weight is set by design rather than fitted towards a more comprehensively measured outcome. The score itself acts to attach a quantity to each trade-off in mutation that this sub-class of silent primer design tools has left implicit; PEEK completes this task in a form the user can personally inspect and re-weight based on the parameters of their own experiment.

Tm additionally holds an extremely important role in primer functionality, as it is the gate that determines which candidate primers are even considered. A systematic Tm model error subsequently doesn’t simply change a number displayed to the user regarding an annealing temperature, but rather it changes the primer sequences the search itself returns. To that end, it became apparent that PEEK’s primer quality rests first on an external check of that calculation, and subsequently its ability to return primers that produce the intended genomic edit requested by the user.

Here, we describe PEEK and test it on two primary grounds. Computationally, we benchmark its Tm and annealing-temperature calculations against the standardized New England Biolabs (NEB) Q5 Tm calculator: the reference published by the manufacturer of the Q5 polymerase whose buffer conditions PEEK functionally models. Specifically, we run all five primer design engines over template sequences that stress the formula’s outputs in order to ensure its accuracy during live experiments. We additionally check each individual silent-mutation proposal suggested by PEEK by re-translating the edited window and searching for the claimed restriction site. Experimentally, we use PEEK to install a KpnI site into the C039_BBz lentiviral chimeric antigen receptor (CAR) plasmid through synonymous substitution and direct insertion; both mutations are later confirmed through nanopore sequencing in order to additionally ensure that no unintended changes exist within the remainder of the 9.9kb construct. Lastly, we test PEEK’s sequence amplification engine against its predicted product size through gel electrophoresis while reporting the limitations that both tests expose.

## MATERIALS AND METHODS

### Implementation

PEEK is written in Python 3.11 and served through Flask, with Biopython handling the sequences. Melting temperatures are computed directly rather than looked up: the base value comes from a closed-form implementation of the SantaLucia (1998) unified nearest-neighbor model (5), and the Owczarzy (2008) treatment (6) then corrects it for the multivalent ion composition of Q5 reaction buffer. Buffer conditions are held at the standard defaults: 500 nM primer, 50 mM monovalent cation, 2.0 mM Mg^2+^, and 0.8 mM total dNTPs. Both the buffer composition and the annealing rule described below are specific to Q5. PEEK’s Tm values are computed under Q5 reaction conditions, and its annealing temperatures follow the convention NEB publishes for that polymerase; neither transfers unchanged to a polymerase supplied with a different buffer or a different annealing convention.

The annealing temperature follows the rule NEB gives for Q5. The rule takes the lower of the two primer Tm values, adds 1 °C, and caps the result at 72 °C, the extension temperature in standard three-step Q5 cycling. We report Tm and annealing temperature separately throughout, and the separation is deliberate. The two quantities behave differently. A melting temperature is a thermodynamic property of a single sequence and is unbounded; a short AT-rich primer may melt below 40 °C, a long GC-rich one above 80 °C. An annealing temperature is a protocol parameter derived from a pair, and it never exceeds 72 °C.

An earlier release (v9) arrived at Tm by a different route. It applied a fitted linear correction, Tm_NEB = 0.585 × Tm_raw + 34.147, to Biopython’s generic nearest-neighbor output, with the coefficients calibrated against a small internal comparison set. A correction of that kind is only defensible inside the range over which it was fitted. Outside that range it is simply wrong, and silently so. We therefore replaced it with the closed-form physical model described above. Whether that replacement was justified is, in part, what the benchmark reported below is meant to test.

### Primer design engines

Five primer design engines share this Tm primitive. The core pair optimizer searches candidate lengths for the forward and reverse primers and scores each pair on the Tm difference between them, on their distance from a 65 °C target and on their combined length. The enzyme-insertion engine appends a restriction site as a 5′ tail and extends both primers to Tm, while enforcing a minimum 15 nt genomic binding tail beyond the inserted site. Two further engines handle placement around a region: the indel/amplification engine sets inward-facing primers flanking a target, whereas the verification engine sets them outside an edited region so that edited and unedited alleles separate as different product sizes on a gel. The site-directed mutagenesis engine is the odd one out. It designs back-to-back overlapping primers and uses a distinct annealing rule, mean Tm minus 3 °C, because those primers anneal to each other rather than independently to the template.

### Silent-mutation search

The silent-mutation engine takes a DNA sequence, which may be a coding sequence or a whole plasmid, together with a target cut position, a position tolerance in base pairs, a codon window size, and an enzyme set. The reading frame is detected rather than supplied. For each of the three frames, the engine measures the stop-codon-free run through the cut position and selects the longest, so an unannotated plasmid can be submitted with only a position nominated. A window of N codons is then centered on that position. Its boundaries are rounded inward onto codon boundaries and verified by assertion, so a proposed mutation cannot split a codon or extend past the end of the sequence, and any window containing an internal stop codon is rejected. Enzymes whose recognition sequence already occurs in the input are removed before the search begins. Any site the engine creates is therefore unique under exact-sequence matching.

Among the codons in the window that have synonyms, the engine enumerates combinations of up to k codon changes (default k = 3), together with every synonymous replacement at each position. Each mutated window is re-translated and discarded unless the amino acid sequence is unchanged. The survivors are then tested against every enzyme in the set for a recognition site whose cut position, taken from that enzyme’s own cut geometry, falls within the requested tolerance. Because the search has to remain interactive in a browser, it is bounded. A preferred enzyme set (SalI, EagI, RsrII, KpnI, MluI, SbfI, NcoI and DpnI) is tried first, the number of variants tested is capped and the search terminates once 50 preferred hits have been found. Each result records the enzyme, the new cut position, the original and mutated window, the translated amino acids, the bases changed and the side of the window on which the site falls.

### Ranking silent-mutation candidates

Surviving variants are ranked by a composite score. It combines host codon usage, two sequence features relevant to translation and the size of the edit itself. A variant that fails a re-check of synonymy, or that contains a stop codon, is assigned a sentinel value of −100 and marked invalid; the ranking therefore cannot return a non-silent edit even if one were somehow to reach this stage. For a valid variant the score, with higher being better, is

`S = w_cai·CAI − w_rare·R − w_struct·F − w_mut·M + w_site·B`

where the five terms and their default weights are those given in Table 1.

**Table 1.**
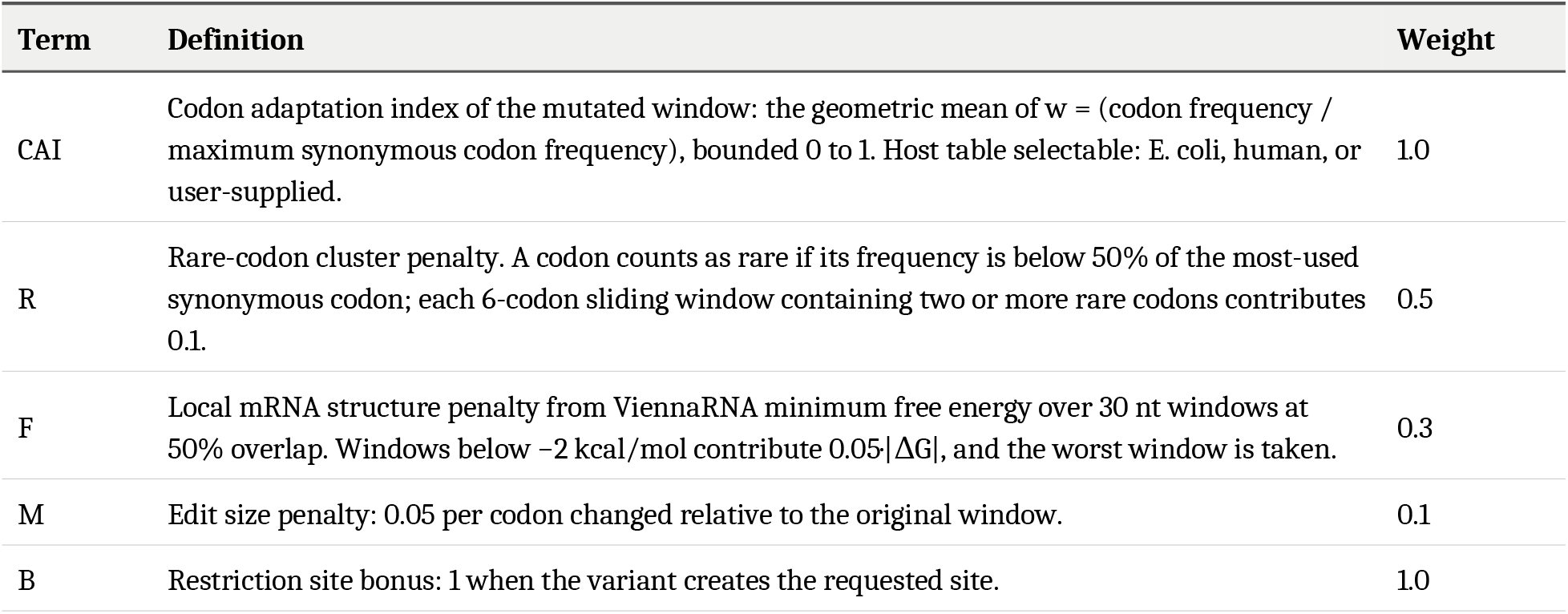
Terms of the composite variant score used to rank candidate silent-mutation edits, with their default weights.

| Term | Definition | Weight |
|---|---|---|
| CAI | Codon adaptation index of the mutated window: the geometric mean of w = (codon frequency / maximum synonymous codon frequency), bounded 0 to 1. Host table selectable: E. coli, human, or user-supplied. | 1.0 |
| R | Rare-codon cluster penalty. A codon counts as rare if its frequency is below 50% of the most-used synonymous codon; each 6-codon sliding window containing two or more rare codons contributes 0.1. | 0.5 |
| F | Local mRNA structure penalty from ViennaRNA minimum free energy over 30 nt windows at 50% overlap. Windows below −2 kcal/mol contribute 0.05·\|ΔG\|, and the worst window is taken. | 0.3 |
| M | Edit size penalty: 0.05 per codon changed relative to the original window. | 0.1 |
| B | Restriction site bonus: 1 when the variant creates the requested site. | 1.0 |

All five terms are returned individually alongside the total, and every weight is user-adjustable, so a ranking can be audited and re-weighted rather than accepted as given. No component is learned or otherwise opaque. The edit-size term M deserves a word of explanation, since it has no counterpart in whole-gene codon optimization, where the sequence is rewritten throughout by design. It is included here because the objective is a different one: to create a requested site with the smallest local change, rather than to maximize expression. Users who care more about which enzyme they obtain than about the expression cost of the edit can instead order candidates by a fixed enzyme preference rank, followed by fewer changed bases.

### Computational benchmarking

The Tm function sits upstream of every design engine, so it was the first thing benchmarked. We did so on sequences chosen independently of PEEK rather than on PEEK’s own output, since the latter would have made the comparison circular. Seventeen synthetic primers (P1–P17) were generated with a fixed random seed to span 15–30 nt and 13–80% GC. These are test sequences and nothing more. They deliberately include short AT-rich oligonucleotides that no engine would propose, so that the benchmark covers the full range over which the function can be called, well beyond the range in which it is normally used.

Each primer was submitted to the NEB Q5 Tm Calculator (7) in batch mode under matching conditions, and the returned value was treated as ground truth. The same primers were combined into nine forward and reverse pairs to test the annealing rule end to end. Two of those pairs were constructed to exceed the 72 °C ceiling, so that the capping behavior could be compared directly.

The silent-mutation engine was tested on a 90 bp coding fragment encoding the first 30 residues of GFP, at five positions spanning the fragment (nt 15, 30, 45, 60 and 75), taking the three highest-ranked proposals at each. Every proposal was re-translated with Biopython outside the engine’s own translation call, and the claimed restriction site was searched for independently in the mutated sequence. Neither check relied on the code being tested.

All five primer design engines were then run against three synthetic templates of 600 bp/35% GC, 800 bp/50% GC and 500 bp/65% GC, chosen to bracket the GC range of typical cloning targets. For each returned primer we checked that it was present in its template, exactly or as the reverse complement, that its individual Tm fell within the 55–72 °C range NEB recommends for Q5 and that the reported annealing temperature was internally consistent with the Tm values in the same result.

### Plasmid editing and sequencing

The C039_BBz plasmid, a 9,944 bp lentiviral CAR construct, was the editing target, with C039_KpnI_RE_site as a positive control already carrying the KpnI site. PEEK’s silent-mutation and enzyme-insertion engines were each asked to install a KpnI recognition sequence at a position within the CD8α coding region. Edits were introduced by whole-plasmid amplification followed by kinase, ligase and DpnI treatment and transformation. Three colonies from the insertion transformation (EI_1, EI_2, EI_3) and one from the silent-mutation transformation (SM_1) were picked and sent for whole-plasmid nanopore sequencing (Plasmidsaurus, order YGH7RT). Separately, the amplification engine was asked for primers spanning a 126 bp region of the same construct; the product was resolved by agarose gel electrophoresis alongside a size ladder. The primer pairs returned by PEEK and used for each of these three reactions are given in Table 2.

**Table 2.**
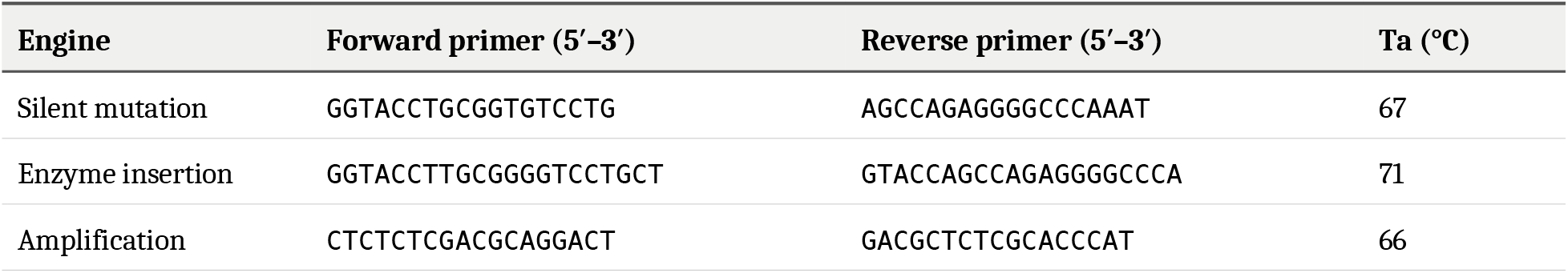
Primer pairs returned by PEEK and used in the experimental validation. The silent-mutation and enzyme-insertion pairs are back-to-back and amplify the whole plasmid; the amplification pair yields a 126 bp product spanning the HIV-1 packaging signal. Annealing temperatures are those reported by PEEK for each pair. Primer lengths, GC contents and binding coordinates, the latter confirmed by exact match against the sequenced assemblies, are given in Supplementary Table S4.

### Sequence analysis

Consensus assemblies were searched for the KpnI recognition sequence GGTACC and then compared base by base against one another across the full plasmid. The point of the full comparison is that any difference outside the intended edit window is detected rather than inferred. The affected coding sequence was translated for the wild-type and edited constructs to test whether the substitutions were silent at the protein level.

The reading frame was established independently rather than assumed. Feature boundaries annotated by pLannotate (8) across the receptor cassette were confirmed to be separated by exact multiples of three, and a contiguous 2,409 bp region spanning five annotated features was translated in the resulting frame. That translation yielded 803 codons with no internal stop. We take this as reasonably strong evidence that the frame used for the silence check is the biological one, since a frameshift would be expected to encounter a stop codon within a few dozen codons of random sequence.

## RESULTS AND DISCUSSION

### Agreement with the NEB Q5 Tm Calculator

Across the 17 primers with varying GC% contents, PEEK’s Tm values agree closely with NEB’s, showing a Pearson R^2^ of 0.9995 and a mean absolute error of 1.32 °C over a 48 °C span, with a root-mean-square error of 1.70 °C and a mean signed bias of +1.16 °C (Figure 1). Computed against the line of exact agreement rather than against a fitted line, R^2^ is 0.982; the difference between the two reflects the small positive bias examined below rather than scatter about the trend. As no fitted correction term is applied anywhere in the calculation, the agreement between the two follows from their shared thermodynamics rather than from fitting PEEK to this comparison set.

**Figure 1.**
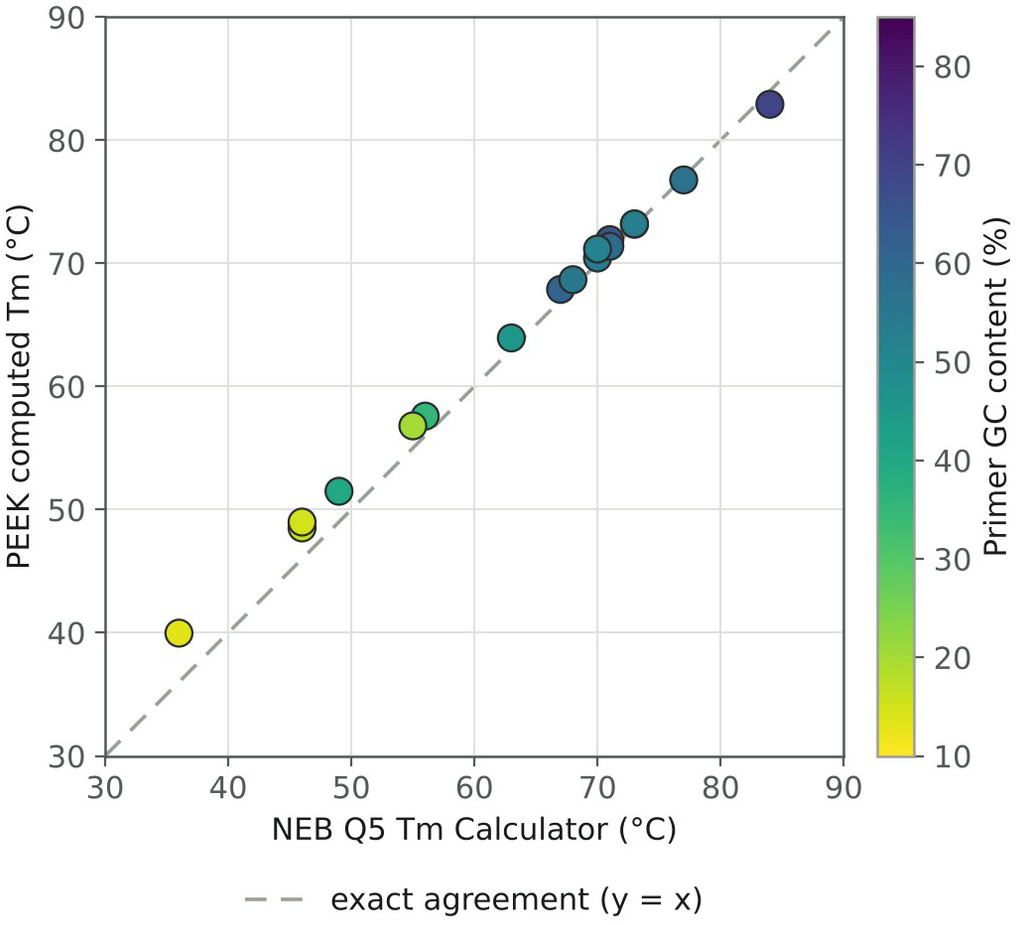
PEEK Tm against NEB Q5 Tm Calculator over 17 synthetic primer test sequences (under matching buffer conditions). Every primer tested falls within 4 °C of exact agreement across a 48 °C range of melting temperatures (Pearson R^2^ = 0.9995; R^2^ = 0.982 against the line of exact agreement; mean absolute error 1.32 °C, root-mean-square error 1.70 °C, n = 17). Individual primer values are given in Supplementary Table S1

The residual of the data is not uniformly distributed. It is largest in the shorter low-GC content primers, falls to near zero around the 50% GC content, and dips slightly negative as the length and GC content incrementally rise (Figure 3). The thermodynamics table in PEEK’s implementation applies a single helix-initiation term (ΔH° = +2300 cal·mol^-1^, ΔS° = +4.1 cal·K^-1^·mol^-1^) once per duplex, whereas the full SantaLucia table distinguishes initiation at terminal GC from AT pairs. Primers that contain terminal AT pairs disproportionately tend to be the short and low-GC content ones where the largest residual is visible.

**Figure 2.**
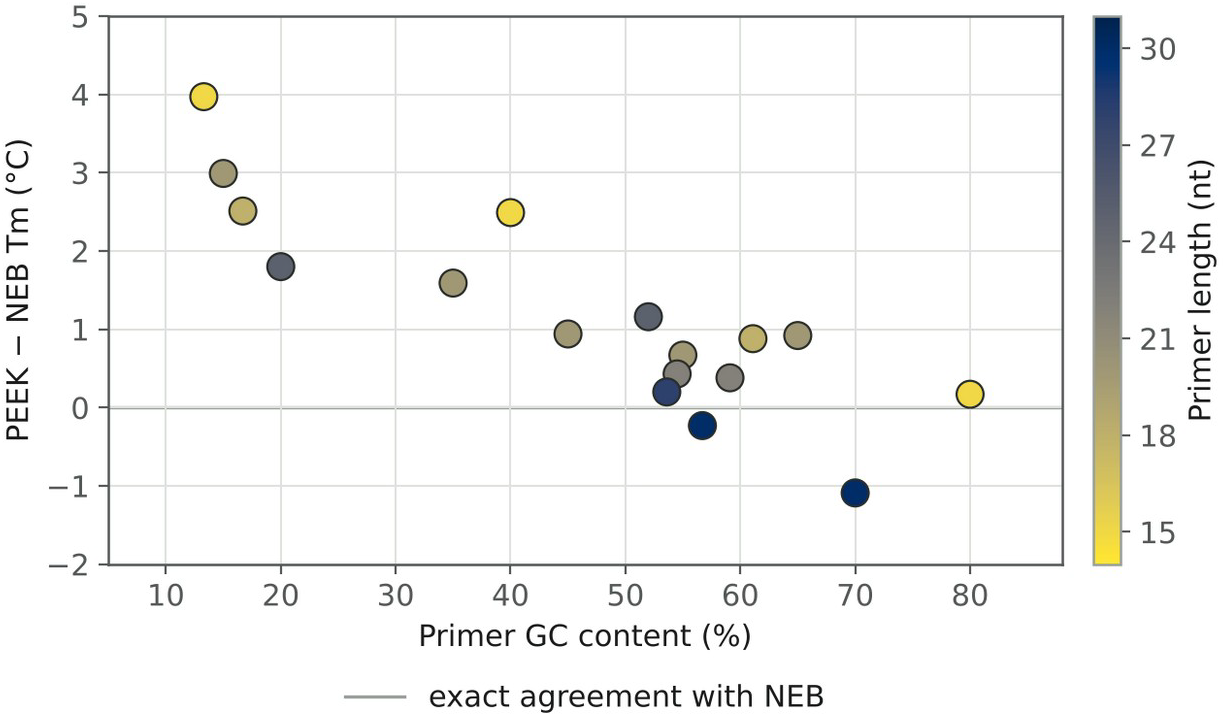
Signed error (PEEK minus NEB) as a function of primer GC content, with point colour encoding primer length. The short, low GC content primers returned the largest residuals. The discrepancy approaches zero for primers of 22 nt or longer, or 50% GC or higher, and becomes slightly negative at the high-GC end.

**Figure 3.**
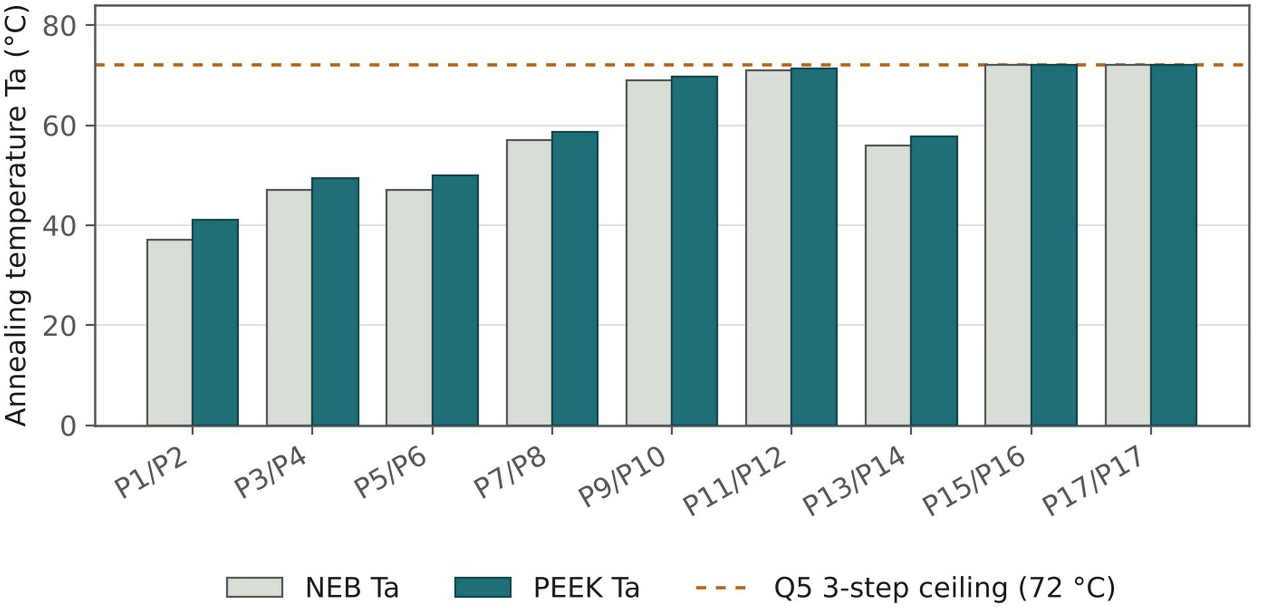
PEEK and NEB computed annealing temperatures for nine primer pairs (drawn from benchmark primers). Unlike the Tm presented in Figure 1, the annealing temperatures in PEEK, likewise to NEB, are bounded by a 72 °C capped temperature. Both primer pairs P15/P16 and P17/P17 were constructed from primers with Tm values that exceed this bound, yet both PEEK and NEB’s calculators properly controlled the Ta to the same value.

This bias was not corrected with some form of fitted offset for three primary reasons. Firstly, NEB reports Tm as an integer, meaning that up to ±0.5 °C disagreement between the two models can be attributed to rounding rather than model error. Secondly, PEEK’s own tolerance for primer pair Tm difference is 5.0 °C, an order of magnitude larger than the bias. Thirdly, the engines target the 60–70 °C range, which typically excludes short, low GC primers where the residual is largest. When primer parameters are restricted to 20+ nucleotides and 35% GC or higher, the median absolute difference from NEB is 0.8 °C. Applying a correction fitted to this comparison set would reintroduce the same failure mode that motivated removing the v9 trendline, namely a correction that holds only across the range it was fitted on. Instead, we record the initiation-term simplification as a candidate refinement.

PEEK’s annealing-temperature rule reproduces NEB’s behaviour exactly, including at the ceiling of 72 °C (Figure 3). Error in this Ta tracks any error in the lower of the two primer Tm values; yet, as the rule implies, Tm values above the ceiling are capped for both NEB and PEEK, and for the two pairs constructed to exceed it the difference collapses to 0.0 °C (72 °C being the extension temperature used in standard three-step Q5 cycling).

### Silent-mutation engine

All 15 primer proposals across the five tested positions preserved the translated protein sequence while simultaneously introducing the intended restriction site, as judged by translation and sequence searches run outside of the PEEK engine. The proposals were designed to introduce one of nine distinct recognition sequences including isoschizomer groups (AluI/AluBI, AgeI/BshTI/AsiGI, DpnI/BstMBI/BssMI, etc), with each proposal requiring between one and four base changes to the construct. We note that this benchmark tests primer correctness rather than completeness as it establishes that the proposals returned by PEEK are valid, not that the genuine finds every possible valid proposal. Doing this would require an exhaustive amount of computational power which we have not performed.

The silent-mutation engine was additionally used to install a KpnI reconstruction site in the C039_BBz construct; the clones resulting from its bacterial transformation were subsequently sequenced. Table 3 provides the individual synonymous codon substitutions performed, with the nucleotide edits being present at the third position of each codon.

**Table 3.** The silent-mutation edits in bacterial clone SM_1 for each codon (54 and 56). Codon numbering in the figure is relative to the start of the CD8α coding fragment within the plasmid. Both substitutions exist at the third codon position and are synonymous, causing no amino acid change in the process. We note that codon 55 which lies between 54 and 56 was also unchanged.

| **Position** | **Codon** | **Codon position** | **Wild-type** | **Edited** | **Wild-type residue** | **Edited residue** |
|---|---|---|---|---|---|---|
| 5244 | 54 | 3rd | ACT | ACC | Thr | Thr |
| 5250 | 56 | 3rd | GGG | GGT | Gly | Gly |

The physical effect this controlled substitution had on the sequence window is shown below. Substituted bases are shown in red, and the KpnI recognition sequence introduced is underlined. Notably, the variant returned by the engine additionally eliminates the only four-guanine nucleotide chain in the window. Because the scoring function of PEEK’s silent-mutation engine penalises local secondary structure, and G-runs contribute strongly to it, the preference for this variant over the single-substitution alternative is consistent with that term operating as intended.

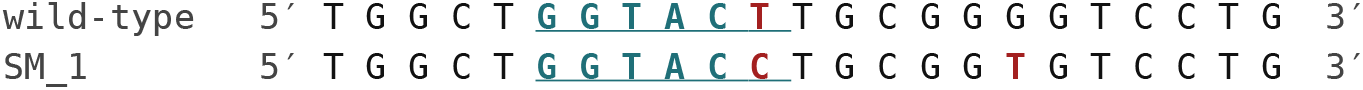
Silent-mutation sequence window spanning between plasmid position 5234 and 5256. The two substitutions occur at position 5244 and 5250. The first substitution completes the KpnI sequence (GGTACC), introducing it into the plasmid synonymously. The second lies downstream of the site and acts to eliminate a four-guanine run present in the parent sequence at 5248 to 5251, the only homopolymer run in this window, while leaving the GC content of the window unchanged.

For further support, translating the affected reading frame returns an identical amino acid sequence both before and after the edit, and across the remaining 9942 bp of the plasmid there exists no difference from the parent construct. Therefore, the edit made was proved to be silent within a sequenced construct rather than a property inferred solely from the primers’ computational design. The translated sequence is given in Supplementary Table S3.

### Tm compliance across the design engines

Of the 30 individual primers tested across PEEK’s five engines, 27 returned Tm’s that fall within the NEB recommended 55-72 °C range, with all 15 primer pairs additionally binding to their source template exactly (Figure 4). The three exceptions to this are not randomly distributed, rather they arise from 2 primary identifiable code paths.

**Figure 4.**
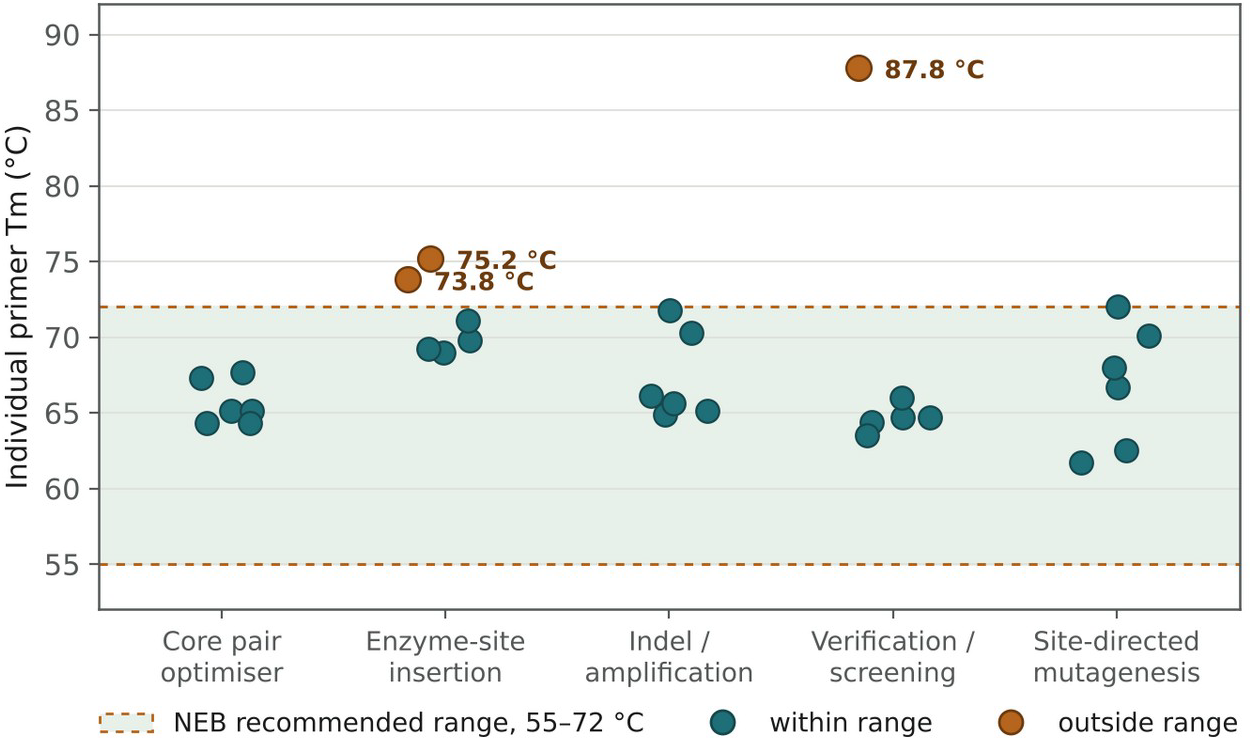
30 individual Primer Tm’s across each design engine. The shaded band marks the NEB recommended Tm range for the Q5 polymerase. Two primer legs from the enzyme-insertion engine and one from the verification engine did not fit into this range. Full engine output is given in Supplementary Table S2.

**Figure 5.**
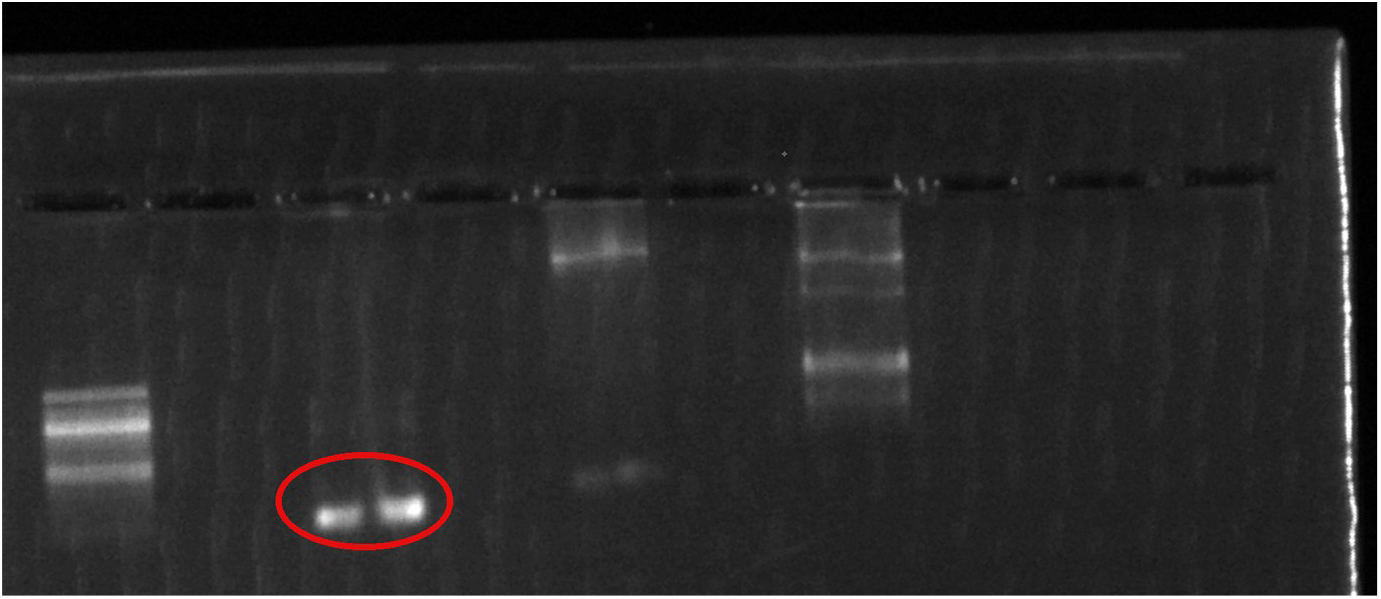
Agarose gel of the product amplified with primers returned by the amplification engine. The circled band is the fastest-migrating species on the gel and sits at the position expected for the predicted 126 bp amplicon relative to the ladder in the adjacent lane.

The two enzyme insertion primers with Tm’s outside the range are forward primers with high GC content. The engine enforces a minimum 15nt length for each individual primer beyond the restriction site to guarantee template contact; thus, when even the minimum length leads to a GC-rich sequence, the Tm can exceed the 72 °C threshold set. This is a deliberate tradeoff which prioritizes the binding of the primer sequence, yet a Tm which breaks the conventional standard should still be treated as an output that needs to be inspected by the user.

The third error is a verification primer with a Tm of 87.8 °C on the 65% GC template. When no candidates between 15-40 nt are deemed to be inside the recommended range, the length of the primer returned is automatically set to the maximum-length (40 nt). A usable primer with an out out-of-range Tm is always more useful to a caller than no primer at all, but the same caveat to the previous errors applies: the user should read the Tm value and judge whether the primer is worth ordering, because the search does not guarantee the value falls in range.

For the core pair, indel/amplification, and site-directed mutagenesis engines no primers with out-of-range Tm’s were produced. This occurred mainly because their algorithms which determine primer length carry neither a fixed-sequence tail nor a fixed anchor with a hard length bound, subsequently reducing the likelihood of a constraint which causes extreme Tm values to be returned.

Regardless, the appropriate approach to the Tm values is to surface the extreme condition to the user explicitly rather than to return an out-of-range primer silently, and we intend to add such a warning.

### Installing a KpnI site in a CAR plasmid

We tested two editing engines and one replication engine on the C039_BBz construct. The target position for restriction site introduction lies within the CD8α coding fragment, between the single-chain variable fragment and the downstream costimulatory and CD3ζ signalling domains, a junction of the kind used for domain-swap experiments and therefore a realistic rather than arbitrary cloning target. The region which was amplified by amplification primers was upstream, however, and is discussed later. All four sequenced samples retrieved from the bacterial cloning resolved as a single intact species of approximately 9.9 kb before sequencing (Figure 6), with per-base read depth at the edit locus between 30× and 209× and no low-confidence base calls in the window.

**Figure 6.**
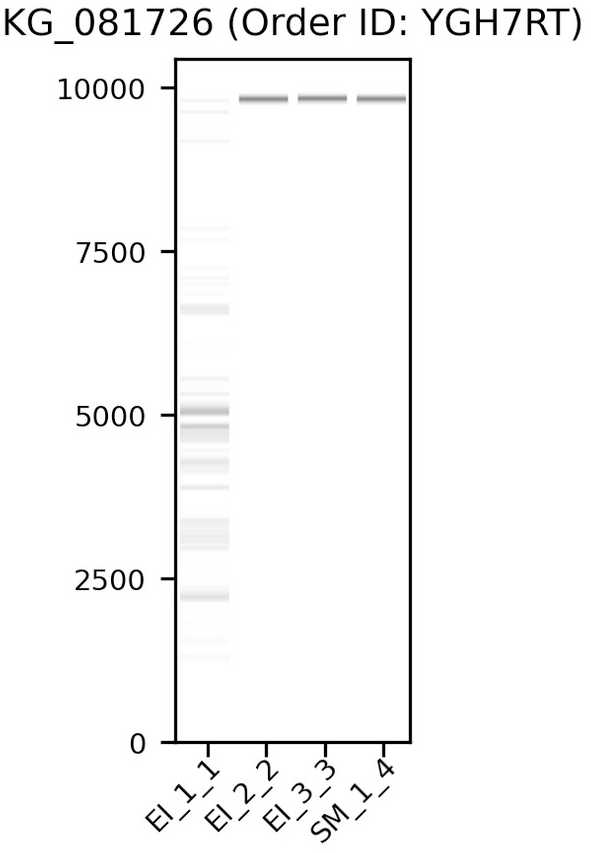
Pre-sequencing fragment analysis of the four submitted clones (Plasmidsaurus order YGH7RT). Each resolves as a single band at approximately 9.9 kb, consistent with intact plasmid and without evidence of degradation or additional species.

### Enzyme-insertion engine

One of the three colonies picked from the “enzyme insertion” bacterial transformation, EI_3, carries a 6 bp GGTACC insertion at the designed junction, lengthening the plasmid from 9,944 bp to 9,950 bp with no other difference anywhere in the construct (Table 4). The remaining two colonies, EI_1 and EI_2, carry no edit and are unchanged from the parent plasmid.

**Table 4.** Whole-plasmid sequencing results for the four sequenced clones. Changes are reported relative to the unedited C039_BBz construct across the complete assembly, not only within the target window.

| **Clone** | **KpnI site** | **Length (bp)** | **Changes relative to parent plasmid** |
|---|---|---|---|
| EI_1 | Absent | 9,944 | None |
| EI_2 | Absent | 9,944 | None |
| EI_3 | Present | 9,950 | 6 bp GGTACC insertion at the designed position, and no others |
| SM_1 | Present | 9,944 | Two synonymous substitutions at the designed positions, and no others |

The three colonies represented by the table are replicate picks from a single transformation plate, not independent design attempts, and the one-in-three recovery rate is a property of the cloning step rather than of the primer. Residual parental plasmid surviving DpnI treatment, and re-ligation without insert both yield unedited colonies at rates that vary between preparations and are independent of primer sequence. The relevant observation for a primer design tool is that where the edit was incorporated, it was incorporated exactly as specified, with no scarring at the junction and no change elsewhere in a 9.9 kb construct. We report the recovery rate because it bears on the experimental effort required, not because it measures design accuracy; distinguishing the two would require a larger panel of clones, which we have not sequenced.

### Amplification engine

The amplification engine returned primers designed to amplify a 126 bp region of the same construct corresponding to the HIV-1 packaging signal (Ψ, positions 1512 to 1637 in the sequenced assembly). Gel electrophoresis of the PCR product resolved a single fast-migrating species whose position relative to the ladder run in the adjacent lane corresponds to the predicted 126 bp amplicon (Figure 5). No additional products were visible in the lane, indicating that the primer pair amplified the single intended target under the conditions used.

### Coordinate conventions

The edit position reported by PEEK (approximately 7665) and the position in the sequenced assembly (approximately 5240) differ because the assembler rotates circular sequences to begin at a recognised landmark. In this case there exists the ColE1 origin, which is not the origin used in PEEK’s reference map. The offset is identical in both edited clones, and the 40 bp of sequence surrounding the edit matches the designed primers exactly in each, confirming that the discrepancy is one of numbering conventions. Typically, edit positions in synthetically engineered plasmids are more robustly reported relative to an annotated feature than as absolute coordinates due to such rotation, and we have adopted that convention here.

### Limitations

The most consequential limitation of this work is what the benchmark does not cover. Here, we have compared PEEK’s Tm model against an external standard (NEB) and its proposals against the constructs they produced in a controlled plasmid, but we have not compared PEEK’s primer selections against those of Primer3, Primer-

BLAST or commercial alternatives on a shared target set, nor its silent-mutation proposals against SDM-Assist (4), which is the closest existing tool and the natural comparator. Such a comparison is the standard against which primer design tools are judged, and until it is performed, our claims are properly read as establishing that PEEK is internally correct, not that it selects better primers or better edits than existing tools.

Several additional limitations are specific to the silent-mutation engine. Its search is heuristic rather than exhaustive: the number of variants tested is capped at 50 preferred hits; as a byproduct, the early stopping point for the primer search could potentially cause a valid edit to be missed, and the absence of a result is not evidence that none exists. Site uniqueness is enforced by exact-sequence matching, so the guarantee holds only for enzymes whose recognition sequence is unambiguous; meanwhile, a degenerate site such as BglI’s GCCNNNNNGGC corresponds to 1024 distinct nucleotide sequences, none of which a literal search for the pattern will find. Within a single run, every returned mutation variant (which corresponds to a primer pair) creates the requested site, so the restriction-site bonus is constant and the ranking is driven entirely by the codon adaptation, rare-codon, structure, and edit-size terms; the bonus only differentiates when scores are compared across enzymes or runs. The default weights are set by design rather than fitted to measured expression or cloning outcomes, which makes the score a transparent prior rather than a validated predictor of protein yield, and we have not tested the expression consequences of any proposed edit to the weights experimentally. Finally, the codon adaptation term is only as meaningful as the host table it is computed against, and the correct table is not always obvious. A lentiviral vector is propagated in E. coli but expresses its insertion in mammalian cells, so an edit scored against the wrong table may be optimised for an organism that never translates the sequence. Built-in tables cover E. coli and humans; any other host would subsequently require a user-supplied table.

The experimental validation rests on a single locus in a single plasmid backbone with a single restriction enzyme, and on one sequenced clone per engine. These results are unambiguous as far as they go, since sequence confirmation across the complete construct leaves little room for interpretation, but they do not establish behaviour across loci of differing GC content, across enzymes with different recognition-sequence composition, or across backbones. The amplification engine was validated by product size against a ladder rather than by sequencing, and the screening and site-directed mutagenesis engines were not tested experimentally outside the synthetic benchmark.

PEEK’s thermodynamic outputs are also scoped to a single polymerase. The primer sequences it returns are not Q5-specific, but the Tm values and annealing temperatures reported alongside them are, and a user working with a different polymerase should treat both as requiring recalculation under that enzyme’s own buffer and annealing rule.

Finally, the computational benchmarks, despite establishing engine correctness, were performed on sequences that were generated rather than encountered. They therefore support the Tm model and the search logic only across the range of conditions tested; moreover, they are a separate line of evidence from the plasmid results and should not be read as extending them.

## CONCLUSION

PEEK’s silent-mutation discovery, restriction-site insertion, amplification, screening, and site-directed mutagenesis primer design are based on a shared thermodynamic basis that agrees with the polymerase manufacturer’s calculator to 1.32 °C mean absolute error across a 48 °C range without any fitted correction. The silent mutation engine creates the smallest synonymous edit at a requested position, and scores candidates with a decomposable objective whose terms are reported individually. The engine was correct on every independently checked proposal. Its proposal held valid when tested physically, with whole-plasmid sequencing confirming two synonymous substitutions at designed positions without changing the rest of the construct. Likewise, the insertion engine produced an exact 6 bp site with no off-target change. Two engines return out-of-range melting temperatures under specific, identifiable conditions, which we document here and intend to surface to users as warnings

The next plausible step is comparison with other tools on the same targets. The results establish that PEEK does what it claims, yet they do not establish how its selections may differ from other kits. That benchmark, paired with more intense wet-lab validation across additional loci, enzymes, and backbones, is the natural next step.

## DATA AVAILABILITY

- **Web server:** https://peekbiology.com. Freely accessible, with no registration required.
- **Source code:** https://github.com/nkura-arch/PEEK-Protein-Engineering-Efficiency-Kit, released under the MIT License.
- **Benchmark data:** the 17 primer sequences with their NEB and PEEK Tm values (Supplementary Table S1), the synthetic templates, the full engine output (Supplementary Table S2), and the translated wild-type and edited reading frames (Supplementary Table S3).
- **Experimental primers:** the three primer pairs used for the silent-mutation, enzyme-insertion and amplification reactions, with their lengths, GC contents, annealing temperatures and binding coordinates (Supplementary Table S4).
- **Plasmid sequencing:** whole-plasmid nanopore sequencing was performed by Plasmidsaurus (order YGH7RT). Consensus assemblies for the four clones reported here are available from the corresponding author on request.
- **Citation:** PEEK is free for academic and non-commercial use. We ask that work using PEEK, in whole or in part, cite this publication.

## ACKNOWLEDGEMENTS

We thank Armin Ghobadi, Bryant Thompson and Oliver Zimmerman for their guidance and support, and the Singh Laboratory at Washington University in St. Louis School of Medicine for providing the plasmid constructs and laboratory facilities used in this work.

